# In vivo Quantitative Tomography of ^225^Ac Daughters in a Prostate Cancer Mouse Model with a Compton Camera

**DOI:** 10.64898/2026.09.18.752779

**Authors:** Biswajit Das, Anil P. Bidkar, S. A. Naik, Victor Negut, David Goodman, M. Streicher, Jaewon Lee, Oscar H. Matousek, Ryan Heller, Megha Basak, Bin Liu, Youngho Seo, Robert R. Flavell, Javier Caravaca

## Abstract

Targeted alpha therapy with ^225^Ac is a promising modality for cancer treatment, as shown in recent preclinical and clinical studies. Preclinical imaging of ^225^Ac is essential to fully understand the biokinetics of developmental ^225^Ac radiopharmaceuticals. Although single photon emission computed tomography (SPECT) is typically used to achieve this in vivo, conventional preclinical scanners struggle to provide high contrast and accurate quantification for ^225^Ac imaging due to the very low injected activities (below 1 kBq*/*g), the unfavorable gamma-ray branching ratios (<26%), and its complex decay chain with several gamma-ray emissions. In this work, we employ a compact CZT-based Compton camera to achieve quantitative tomography of ^225^Ac daughters in live mice at 18.5 kBq (0.5 *µ*Ci) injected activities. Three mice bearing 22Rv1 prostate cancer subcutaneous xenografts were scanned in a single bed position for 30 minutes following intravenous administration of 18.5 kBq of the Anti-CD46 ^225^Ac-Macropa-PEG_4_-YS5 conjugate. We successfully visualized the in vivo biodistributions of the ^225^Ac daughters ^221^Fr and ^213^Bi, distinguishing between the tumor xenograft and the central organs. Quantification of the tumor uptake from the images revealed an activity as low as 1.1 kBq. In vivo activity quantification computed from the images was compared to ex vivo biodistribution measurements using a gamma counter after harvesting the tissue, showing strong agreement.

## 1. Introduction

Targeted alpha therapy (TAT) has emerged as a highly promising modality for the treatment of both solid and hematologic cancers, owing to the unique radiobiological properties of alpha particles (*1*–*4*). Their short path length (50–100 *µ*m) and high linear energy transfer (~100 keV/*µ*m) enable highly localized energy deposition, resulting in dense DNA double-strand breaks and potent cytotoxicity while sparing surrounding healthy tissue (*5*). Among available alpha emitters, actinium-225 (^225^Ac) is regarded as a promising isotope for TAT (*6*–*13*), due to its relatively long 9.9-day half-life and decay chain that yields four main alpha emissions (*1*). A schematic representation of the decay chain of ^225^Ac and the ^225^Ac-based TAT is shown in Fig. 1.

**Figure 1.**
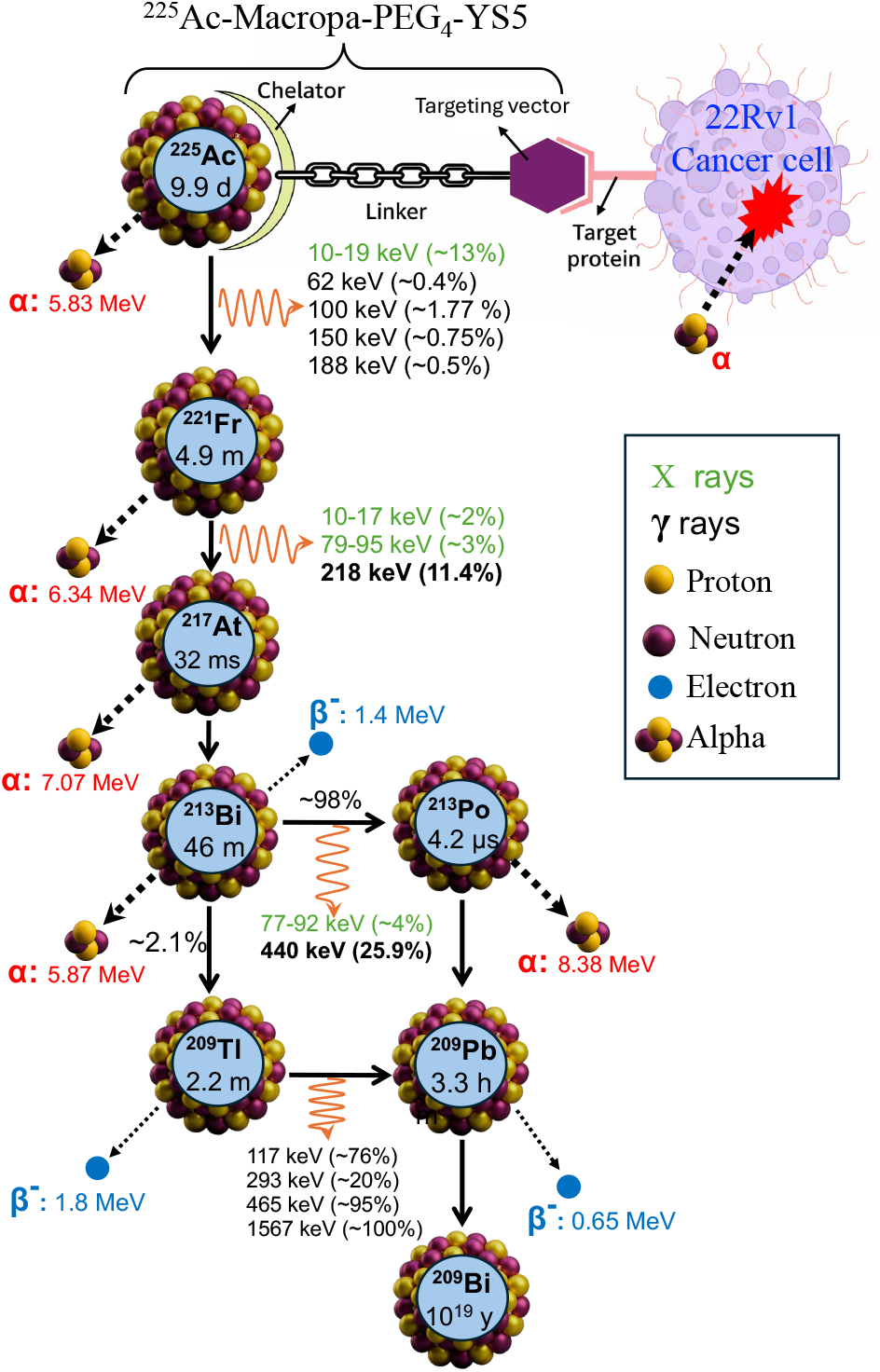
Schematic of the ^225^Ac-Macropa-PEG_4_-YS5 radiopharmaceutical in targeted alpha therapy and the corresponding ^225^Ac decay chain, showing the *α*- and *β*-emitting daughter radionuclides, the key gamma emissions at 218 keV (^221^Fr, 11.4%) and 440 keV (^213^Bi, 25.9%), and the decay to stable ^209^Bi.

Despite its promise, the development and optimization of novel ^225^Ac-based radiopharmaceuticals remain hindered by challenges in accurately assessing in vivo biodistribution and dosimetry. Following alpha decay, daughter nuclides may recoil and dissociate from the parent compound, leading to off-target accumulation and unintended radiation dose to healthy tissues (*14*–*16*). This phenomenon complicates toxicity assessment and underscores the need for reliable methods to track both the parent radionuclide and its progeny in vivo. Currently, evaluation of therapeutic efficacy and toxicity largely relies on ex vivo biodistribution studies, which require sacrificing multiple animals, harvesting their organs, and quantifying activity using gamma or alpha counters to determine organ-specific dose as a function of treatment time points (*17*–*19*).

Noninvasive in vivo imaging of ^225^Ac and its daughter nuclides would enable longitudinal pharmacokinetic assessment, 3D dose mapping, and improved understanding of radionuclide redistribution during therapy. Alpha and beta particles are not suitable for external detection due to their limited penetration in tissue, and alternative approaches such as Cherenkov imaging suffer from poor spatial resolution and lack radionuclide specificity (*20*).

The preferred approach for imaging the biodistribution of ^225^Ac daughters is to detect the gamma rays emitted following their alpha/beta decays. However, this is extremely challenging because, due to the potency of alpha particles, very low activities below 1 kBq*/*g are typically administered in mice. Additional limitations arise from the low branching ratios of the two dominant gamma-ray emissions, 440 keV from ^213^Bi and 218 keV from ^221^Fr, which are approximately 26% and 11.4%, respectively (*21*), further limiting the feasibility of in vivo imaging.

In-vitro imaging of ^225^Ac daughter nuclides has been demonstrated using a commercial preclinical micro SPECT/CT platform (*22*). However, this required multi-hour acquisition times and phantom activity levels approximately two orders of magnitude above acceptable toxicity limits, thereby precluding practical in vivo imaging. Very recently, a cadmium zinc telluride (CZT)-based SPECT with a novel collimator design, the Alpha-SPECT-Mini, was developed for preclinical imaging of beta and alpha emitters (*23*, *24*) and evaluated with high ^225^Ac concentrations of 13.7 MBq*/*mL in phantoms scanned for several hours in several bed positions (*25*).

Compton imaging has been explored in the past years by us and others as an alternative to preclinical SPECT. A simulation study of a Compton camera incorporating GAGG crystals showed an encouraging performance in virtual phantoms (*26*). Nevertheless, the reported results relied on activity levels substantially greater than (3-8 MBq/mL) those typically administered in preclinical experiments. Recently, a dual modality imaging system consisting of two high-purity germanium double-sided strip detectors that can be used as a Compton camera or a coded aperture camera was reported for ex vivo mouse imaging of ^225^Ac daughter biodistribution, requiring acquisition times of several hours (*27*).

In our previous simulation studies (*28*), we demonstrated that Compton imaging with a CZT camera is a highly promising approach for in vivo imaging of ^225^Ac daughter nuclides, owing to its collimator-free design and inherently higher sensitivity compared with conventional collimated systems. In our recent work, we proposed a single-layered, 3D-positioning, CZT-based Compton camera in order to achieve higher sensitivity for tomography of ^225^Ac daughters and evaluated the approach in experiments using a mouse phantom (*29*), concluding that using this camera at a single bed position, about 23 minutes scan would be required to obtain images of sufficient quality for tumor visualization in an in vivo scenario.

In this study, we obtain in vivo 3D images of ^225^Ac daughters (^213^Bi, and ^221^Fr) in mice at therapeutically relevant activities with injections below 1 kBq*/*g. Scanning the mice in a single bed position for 30 minutes with a system with only one Compton camera head, we visualized the in vivo biodistribution of ^221^Fr and ^213^Bi. Imaging was performed in a prostate cancer model following administration of an ^225^Aclabeled anti-CD46 human antibody binding to a tumor-selective epitope (*30*), enabling visualization of tumors and major organs. More importantly, quantitative imaging results showed strong agreement with ex vivo biodistribution measurements. To the best of our knowledge, this study represents the first demonstration of in vivo 3D imaging of the most relevant ^225^Ac daughter radionuclides in mice at therapeutically relevant activity levels below 1 kBq*/*g.

## 2. Approach

### 2.1. Compton imaging and image reconstruction

In vivo imaging was performed using a single-layer 3D positioning CZT Compton camera (M400, H3D Inc., USA), previously characterized for ultra-low-activity radionuclide imaging (*29*). The experimental setup is shown in Fig. 2(a). Two-interaction (*nHit* = 2) Compton events corresponding to the 218 keV emission of ^221^Fr and the 440 keV emission of ^213^Bi were selected for image reconstruction. A schematic representation of the Compton imaging process for in vivo mouse imaging through our system is shown in Fig. 2(b). Images were reconstructed using POSSUM, a Python-based GPU-accelerated list-mode Ordered Subset Expectation Maximization (OSEM) framework for Compton image reconstruction recently developed by our group (*31*), using detector-specific angular response parameters and 50 iterations and one subset. Detailed descriptions of the imaging system, event selection, in vivo image reconstruction, ROI calibration, ex vivo biodistribution measurements, and activity recovery analysis procedure are provided in the Supplemental Material (Sec. I)

**Figure 2.**
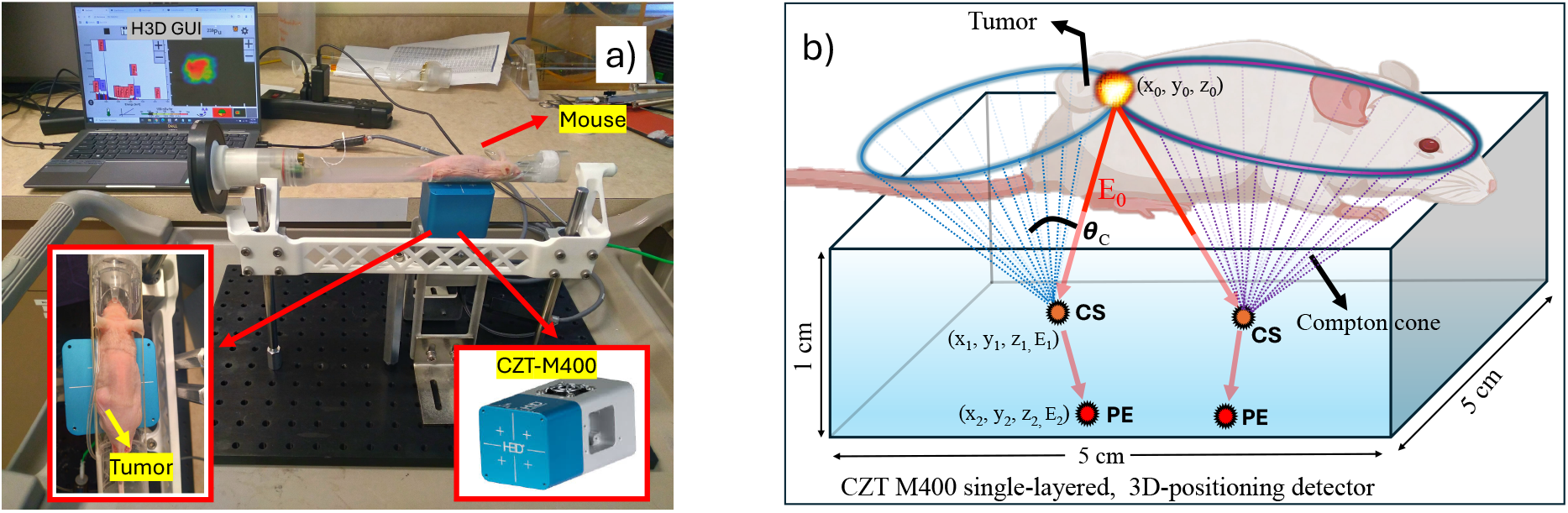
a) Experimental setup showing one of the mice on the mouse bed scanned by the M400 Compton camera connected to the data acquisition computer. The insets show the upper view of the mouse (left) and the M400 detector (right). b) Schematic of the Compton imaging process for in vivo mouse imaging. Gamma rays of energy *E*_0_ undergo Compton scattering (CS) followed by photoelectric absorption (PE), making Compton cones from measured energies (*E*_1_, *E*_2_) and interaction positions. The collective intersections of multiple Compton cones are used to reconstruct the 3D distribution of the radioactive source.

### 2.2. Prostate cancer mouse model and ^225^Ac therapy

We used prostate cancer tumor models developed in six-week old male athymic mice (n=3). Approximately 2 *×* 10^6^ 22Rv1 prostate cancer cells were implanted into the nude animals with Matrigel (50:50). Three weeks after cell implantation, once tumor growth was established, the mice were injected with 0.5 *µ*Ci (18.5 kBq) of ^225^Ac-Macropa-PEG_4_-YS5 radiopharmaceutical (*32*) via the tail vein. A schematic of the ^225^Ac-Macropa-PEG_4_-YS5 radiopharmaceutical, together with the associated *α*-, *β*-, and *γ*-emissions throughout the ^225^Ac decay chain for TAT, is shown in Fig. 1. On the seventh day following radiopharmaceutical administration, the mice were anesthetized, and in vivo imaging scans were performed using our imaging system (Fig. 2(a)).

### 2.3. Imaging experiment

Experimental data were acquired in list-mode format using the web-based graphical user interface provided by H3D Inc. and stored in binary files. Gamma-ray energy calibration was performed using the vendor-provided software with ^57^Co and ^137^Cs sources, and sub-pixel position calibration preformed using ^57^Co and ^225^Ac sources, with gamma-ray energies of 122 keV, 218 keV, and 440 keV. For each mouse, a 30-minute data acquisition was performed using a single-bed position, as shown in Fig. 2(a). Gamma-ray emissions originating from ^225^Ac and its daughter radionuclides (shown in Fig. 1) were recorded. These data are converted into a list-mode CSV file fed to the image reconstruction algorithm (see Supp. Mat. Sec. I.C). Following gamma-ray imaging scans, the mice were transported in the same mouse bed to a commercial preclinical *µ*SPECT/CT system (VECTor4/CT (*33*)) where computed tomography (CT) scans were acquired for subsequent anatomical coregistration.

## 3. Results

### 3.1. ^225^Ac gamma-ray spectrum in tumor-bearing mice

Gamma rays emitted from ^225^Ac and its daughter nuclides distributed across various organs in the mice were detected using our imaging system. The energy spectrum obtained from one of the mice is shown in Fig. 3(a). Within the energy range of 30–500 keV, most of the characteristic gamma rays and X-rays emitted by the decay chain of ^225^Ac, including emissions from ^225^Ac (62 keV, 100 keV, 150 keV, and 188 keV), ^221^Fr (218 keV), ^213^Bi (440 keV), and ^209^Tl (117 keV, 293 keV, and 465 keV) are clearly identified. The energy resolutions at 218 keV and 440 keV were approximately 1.3% and 0.7%, respectively. At energies below 100 keV, several low-intensity *X*-ray and low-energy gamma-ray emissions are present, which are largely attenuated in the detector entrance window and tend to merge in the measured spectrum.

**Figure 3.**
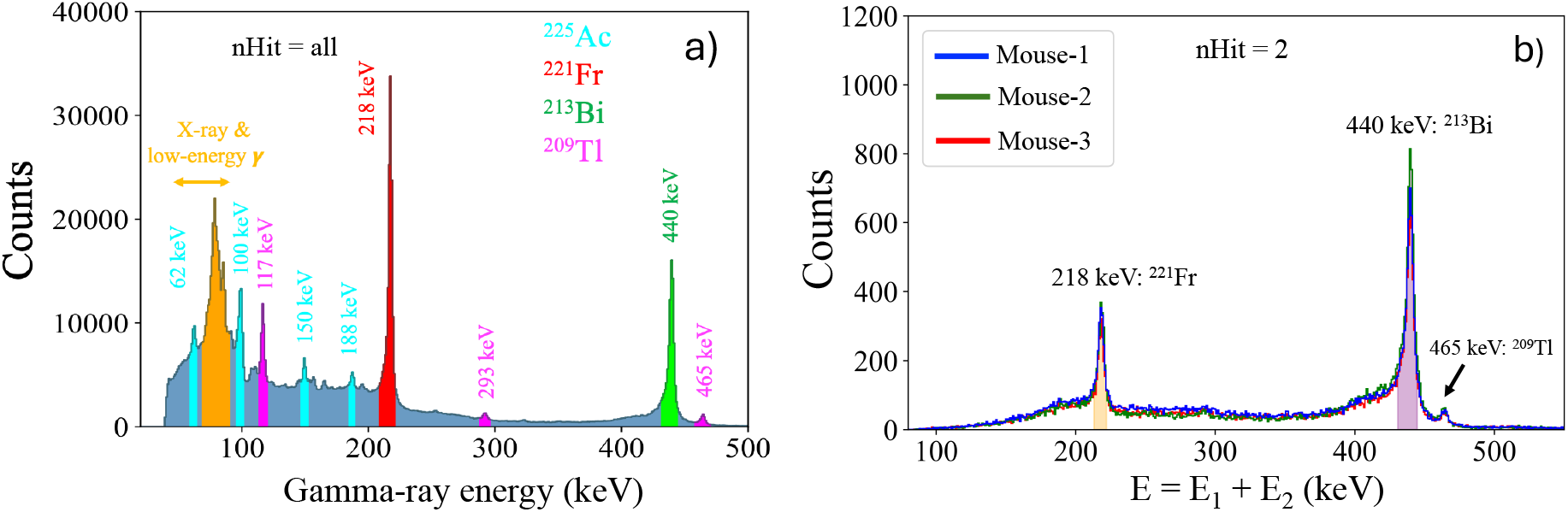
a) Measured gamma-ray energy spectrum of ^225^Ac and its daughter nuclides in one mouse, showing characteristic emissions within 30–500 keV, including contributions from ^225^Ac, ^221^Fr, ^213^Bi, and ^209^Tl. b) Summed energy spectrum (*E*_1_ + *E*_2_) for *nHit* = 2 events of three mice. Prominent summed-energy peaks, highlighted at 218 keV (^221^Fr) and 440 keV (^213^Bi), are observed and correspond to the principal gamma-ray emissions in the ^225^Ac decay chain used for image reconstruction.

### 3.2. Energy-summed spectrum analysis

The measured summed-energy spectrum (*E*_1_ + *E*_2_) for two-interaction (*nHit* = 2) events from the three mice is shown in Fig. 3(b). Prominent peaks corresponding to 218 keV from ^221^Fr and 440 keV from ^213^Bi are clearly observed, reflecting the dominant gamma-ray emissions from the ^225^Ac decay chain suitable for Compton reconstruction. In contrast, the 465 keV emission from ^209^Tl appears with low intensity, and other low-energy gamma rays and *X*-rays are not distinctly visible in the summed spectrum, due to lower branching ratios and lower Compton scattering cross-sections at lower energies. Therefore, Compton imaging of those emissions are beyond the scope of this study.

### 3.3. In vivo imaging of ^225^Ac daughters

The two principal daughter isotopes, ^213^Bi and ^221^Fr, were imaged in vivo using Compton imaging in 30-minutes scan by selecting the 218 keV and 440 keV peaks in Fig. 3(b). The experimental imaging results for these two daughters are presented in the following subsections.

#### 3.3.1 ^213^Bi Compton imaging

The maximum intensity projections (MIP) of Compton images of ^213^Bi daughter nuclides of three mice, overlaid on the corresponding CT images, are shown in Fig. 4(a). The coronal, transverse, and sagittal slices through the tumor of the CT and fused CT + 440 keV Compton images for the three mice are shown in Fig. 4(b–d). Tumor images are clearly visualized in the Compton images of all three mice. The corresponding tumor uptake values (in nCi) are summarized in Table 1. The spatial correspondence between the Compton signal and the tumor location on CT demonstrates successful targeting and retention of the radiopharmaceutical. However, the tumor signal is small in mouse-3 (Fig. 4(d)) because of its smaller tumor size. Uptake in major central organs (liver, heart, kidneys, and lungs) appear merged in a single spot in the reconstructed images. This effect can be attributed to the limited angular resolution of the imaging system, as well as insufficient counting statistics to resolve closely spaced structures with distinct activity distributions. Despite this limitation, the overall central uptake region remains identifiable, providing a qualitative representation of the non-tumor uptake. Quantitative analysis of the reconstructed images was performed using nCi-level source calibration, as described in Sec. 3.4.

**Table 1:**
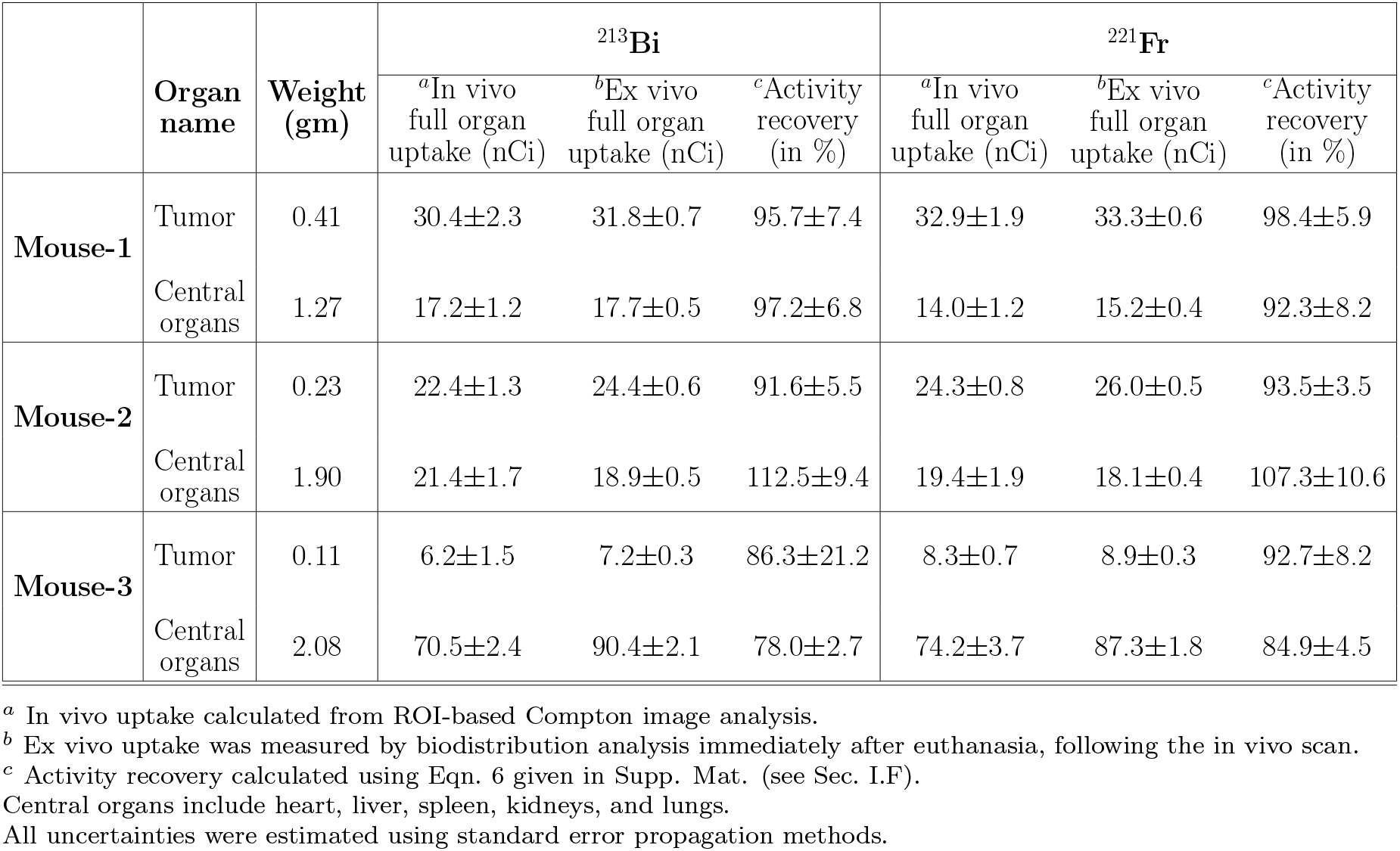
Quantitative comparison between in vivo Compton imaging and ex vivo biodistribution measurements for ^213^Bi and ^221^Fr, in three mice. The table summarizes the quantified uptake activities (nCi) in the tumor and combined central organ regions obtained from ROI-based in vivo image analysis and the corresponding ex vivo measurements, together with the calculated activity recovery values.

| | Organ name | Weight (gm) | $^{213}\text{Bi}$ | | | $^{221}\text{Fr}$ | | |
| --- | --- | --- | --- | --- | --- | --- | --- | --- |
|  |  |  | <sup>a</sup>In vivo full organ uptake (nCi) | <sup>b</sup>Ex vivo full organ uptake (nCi) | <sup>c</sup>Activity recovery (in %) | <sup>a</sup>In vivo full organ uptake (nCi) | <sup>b</sup>Ex vivo full organ uptake (nCi) | <sup>c</sup>Activity recovery (in %) |
| Mouse-1 | Tumor | 0.41 | 30.4±2.3 | 31.8±0.7 | 95.7±7.4 | 32.9±1.9 | 33.3±0.6 | 98.4±5.9 |
| | Central organs | 1.27 | 17.2±1.2 | 17.7±0.5 | 97.2±6.8 | 14.0±1.2 | 15.2±0.4 | 92.3±8.2 |
| Mouse-2 | Tumor | 0.23 | 22.4±1.3 | 24.4±0.6 | 91.6±5.5 | 24.3±0.8 | 26.0±0.5 | 93.5±3.5 |
| | Central organs | 1.90 | 21.4±1.7 | 18.9±0.5 | 112.5±9.4 | 19.4±1.9 | 18.1±0.4 | 107.3±10.6 |
| Mouse-3 | Tumor | 0.11 | 6.2±1.5 | 7.2±0.3 | 86.3±21.2 | 8.3±0.7 | 8.9±0.3 | 92.7±8.2 |
| | Central organs | 2.08 | 70.5±2.4 | 90.4±2.1 | 78.0±2.7 | 74.2±3.7 | 87.3±1.8 | 84.9±4.5 |

<sup>a</sup> In vivo uptake calculated from ROI-based Compton image analysis.
<sup>b</sup> Ex vivo uptake was measured by biodistribution analysis immediately after euthanasia, following the in vivo scan.
<sup>c</sup> Activity recovery calculated using Eqn. 6 given in Supp. Mat. (see Sec. I.F).
Central organs include heart, liver, spleen, kidneys, and lungs.
All uncertainties were estimated using standard error propagation methods.

**Figure 4.**
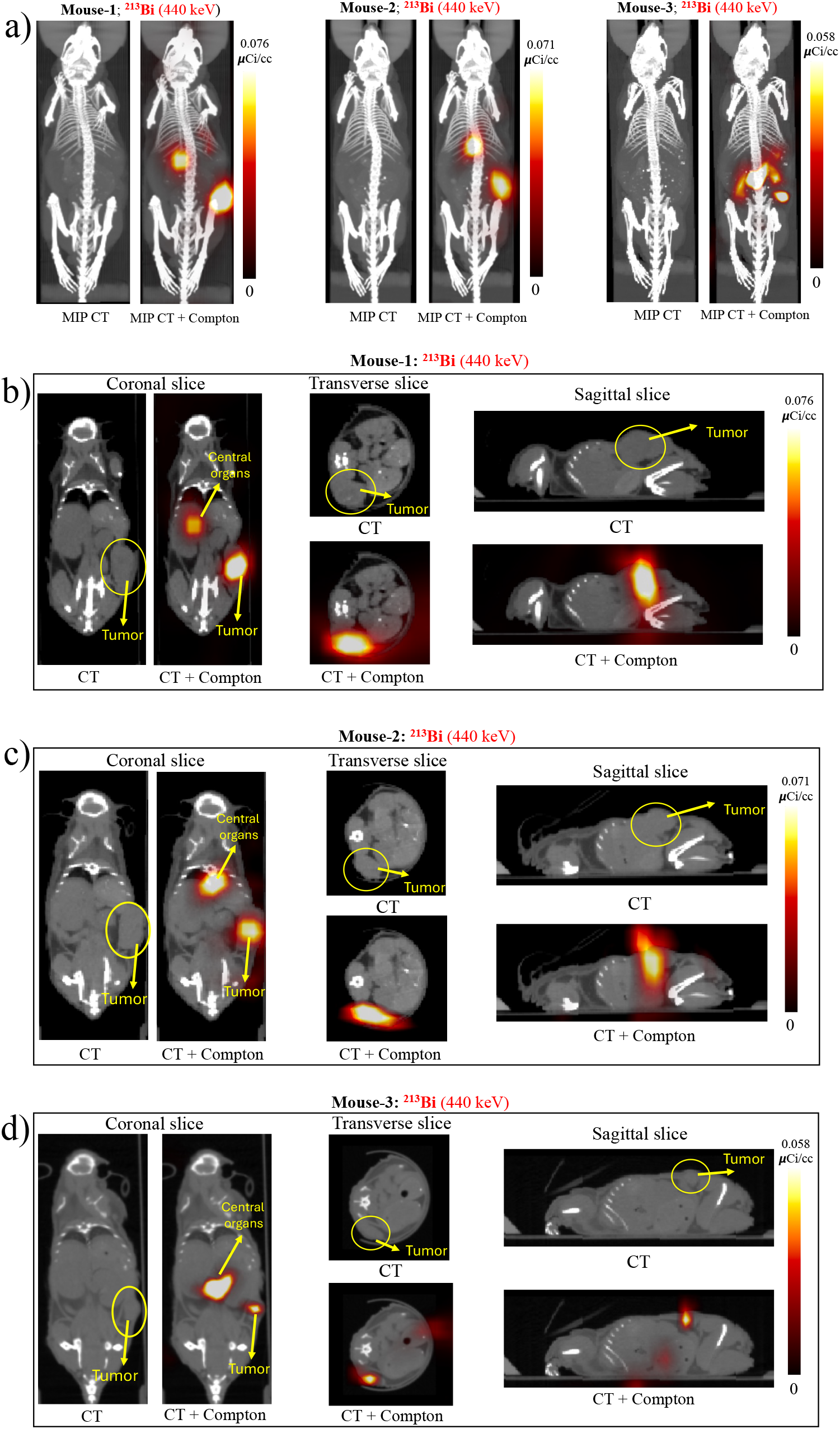
Coregistered Compton and CT images of ^213^Bi in three tumor-bearing mice, acquired at a single bed position with a 30-min acquisition. (a) Maximum-intensity projections (MIPs) of the reconstructed CT and fused CT + Compton images (440 keV; ^213^Bi) for the three mice. (b–d) Coronal, transverse, and sagittal slices through the tumor from the reconstructed CT and fused CT + 440 keV Compton images in three mice. Tumors are clearly visualized in all three mice.

#### 3.3.2. ^221^Fr Compton imaging

The maximum intensity projections of the reconstructed ^221^Fr Compton images, overlaid on the corresponding CT images for the three mice, are shown in Fig. 5(a). Coronal, transverse, and sagittal slices through the tumor from the CT and fused CT + 440 keV Compton images of three mice are presented in Fig. 5(b–d). Tumor uptake of ^221^Fr is clearly visualized in the Compton images for all three mice. The reconstructed ^221^Fr distributions show uptake patterns broadly similar with those observed for ^213^Bi, with activity localized in the tumor and central body region. In all mice, the tumors remain discernible in Fig. 5(b–d), confirming that the 218 keV emission can also be used to visualize tumor-associated activity following administration of the ^225^Ac-labeled radiopharmaceutical through Compton imaging.

**Figure 5.**
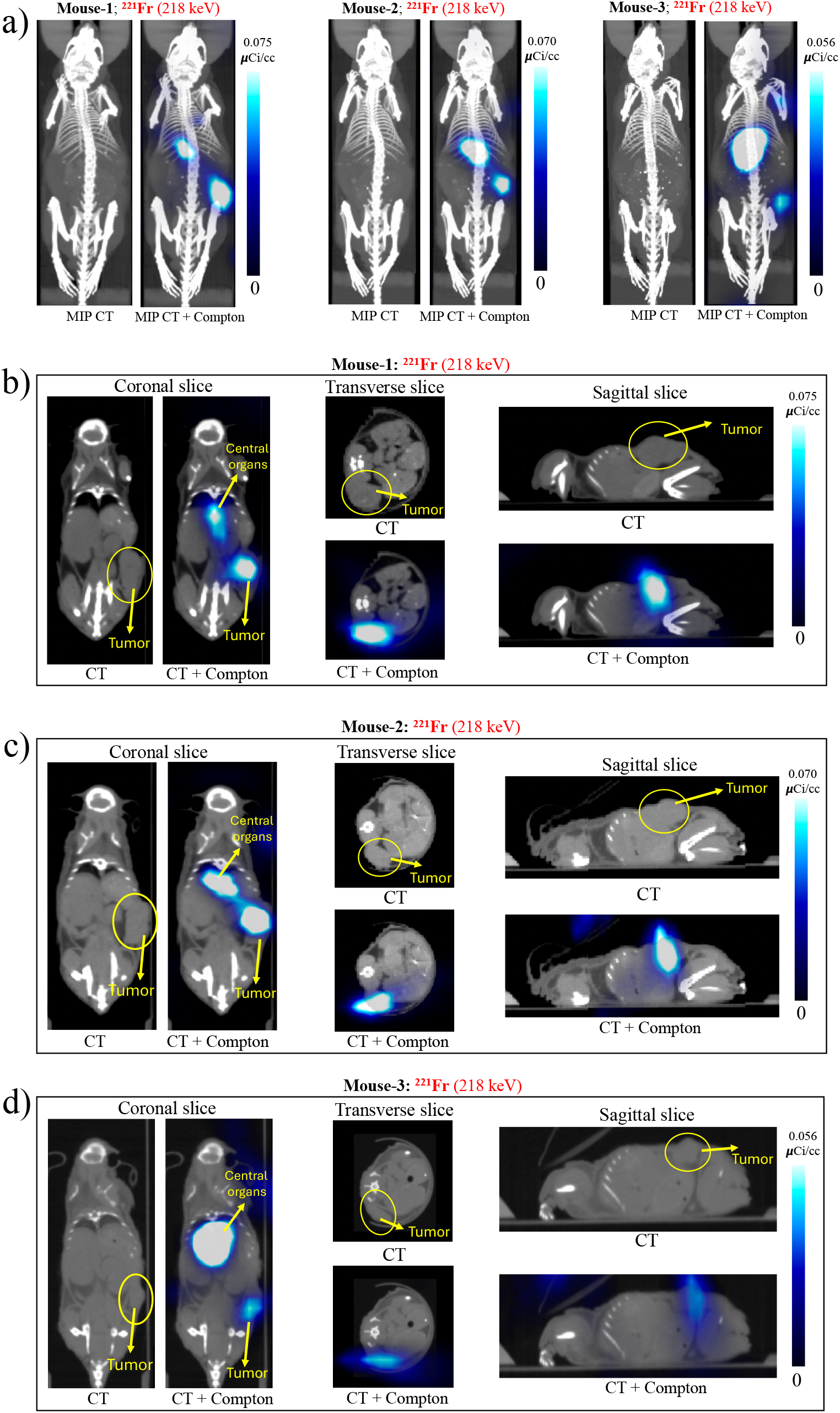
Compton images of ^221^Fr acquired in three tumor-bearing mice using a single-detector configuration (30-min acquisition, single bed position) were coregistered with the corresponding CT images.(a) The MIP of the reconstructed CT and fused CT + Compton images (218 keV; ^221^Fr) for the three mice. (b–d) Coronal, transverse, and sagittal slices through the tumor from the reconstructed CT and fused CT + 218 keV Compton images in three mice. Tumors are clearly visualized in all three mice.

Here also, in mouse-3, the tumor signal is slightly weak (Fig. 5(d)), which is consistent with its smaller tumor burden and lower localized uptake (also presented later in the Table 1). Compared with the ^213^Bi images, the ^221^Fr images exhibit slightly spatial blurring, which is expected from the broader ARM at 218 keV and the correspondingly lower reconstruction precision. As in the ^213^Bi images, uptake in the major central organs (liver, heart, kidneys, and lungs) appears merged into a single region of activity but remains identifiable.

### 3.4. In vivo image quantification

The ROI counts obtained within the 440 keV (^213^Bi) and 218 keV (^221^Fr) energy windows for the point sources were plotted as a function of source activity, as shown in Fig. 6(a,b). Both datasets (^213^Bi and ^221^Fr) exhibited strong linearity, with coefficients of determination (*R*^2^) approaching unity (*≈* 1), indicating a proportional relationship between detected counts and activity over the investigated range.

**Figure 6.**
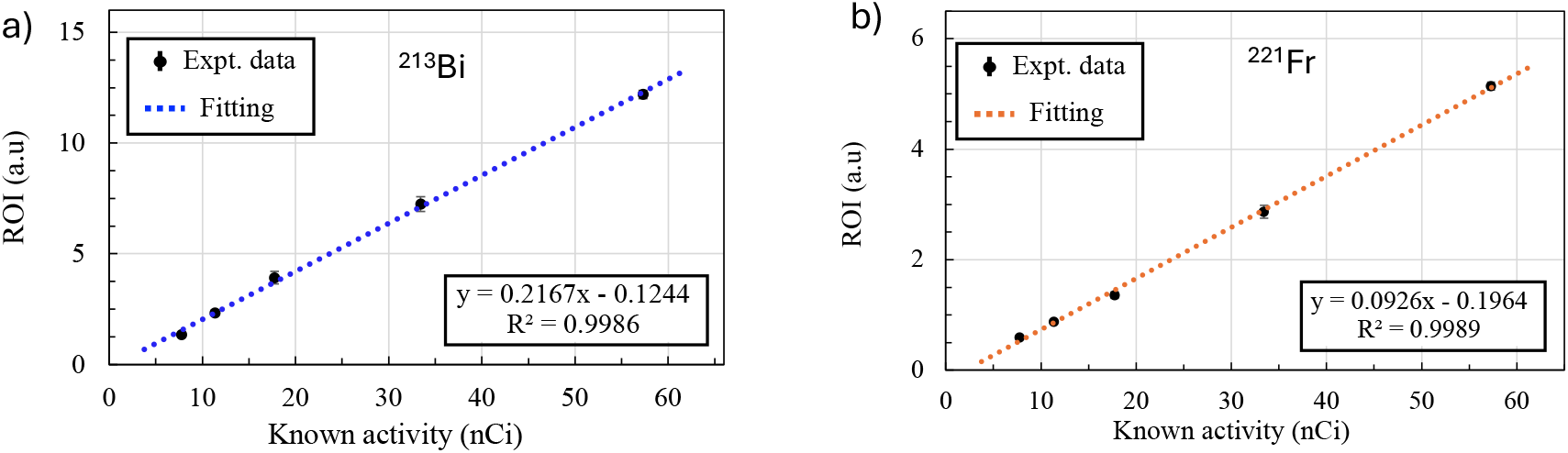
Calibration curves used for activity quantification. Total ROI counts obtained from reconstructed 3D Compton images of ^225^Ac point sources are plotted as a function of source activity for (a) the 440 keV peak (^213^Bi) and (b) the 218 keV peak (^221^Fr).

The quantified activities (nCi) for the tumor and central activity regions in all three mice, determined as described in the Supp. Mat. (Sec. I.D), are summarized in Table 1. These quantified activity values were also compared with the ex vivo biodistribution measurements (also given in Table 1) and showed good agreement, as described in the following section.

### 3.5. Ex vivo biodistribution experimental results

From the ex vivo biodistribution measurements (procedure described in Supp. Mat. Sec. I.E), the total uptake activities of ^213^Bi and ^221^Fr in the tumor and the combined central organ region were determined for all mice and expressed in nCi. These results along with the measured mass of each specimen are summarized in Table 1 alongside the corresponding in vivo quantification results. The measured mass of each specimen and the corresponding ex vivo activity concentrations (%/gm) for the tumor and central organs region are also provided in the Supp. Mat. (Sec. II.A).

### 3.6. Activity recovery

The activity recovery values (procedure described in Supp. Mat. Sec. I.F) for the ^213^Bi and ^221^Fr daughter nuclides in tumors and combined central organ regions, aggregated across all three mice, are presented in Fig. 7(a,b). For tumor regions, both ^213^Bi and ^221^Fr images demonstrate good activity recovery for all mice, typically in the range of 86%–98%, as summarized in Table 1. These results indicate reliable quantitative performance of the Compton imaging approach for localized activity distributions. For both ^213^Bi and ^221^Fr, the uncertainty ranged from approximately 3% to 10%.

**Figure 7.**
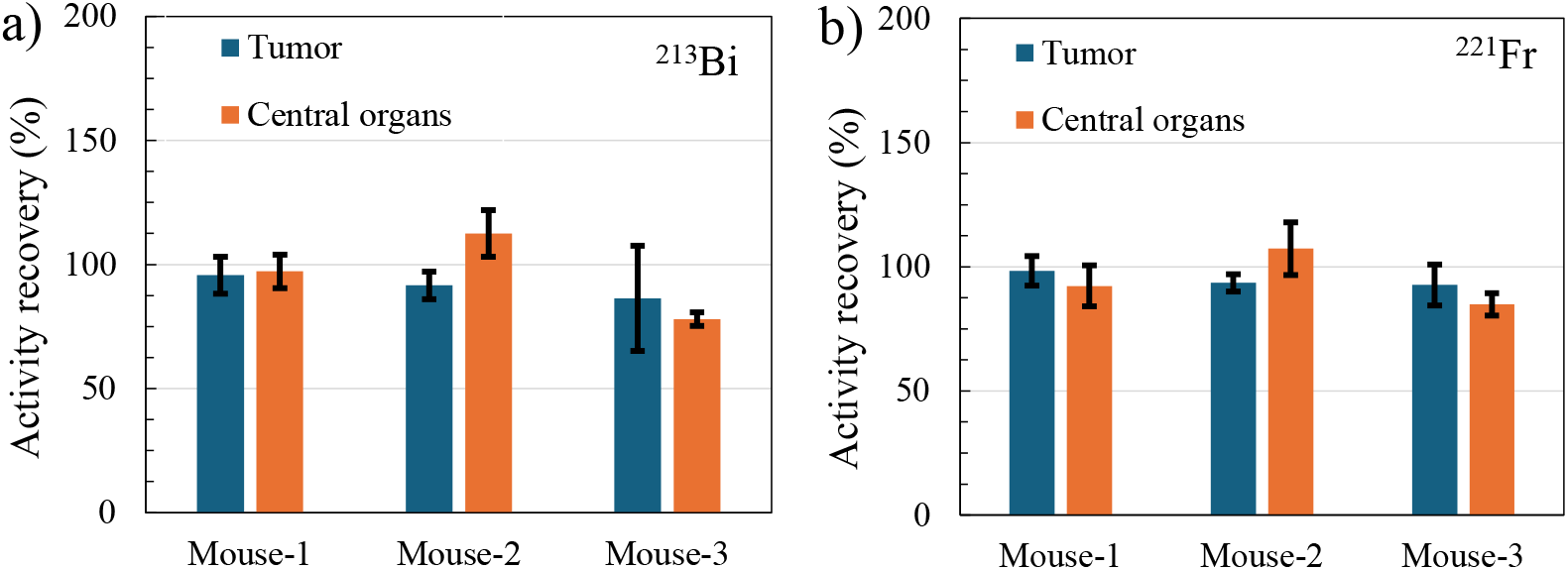
Activity recovery for (a) ^213^Bi and (b) ^221^Fr daughters of tumor and aggregated central organs derived from ROI-based in vivo quantification and ex vivo measurement.

## 4. Discussion

This study demonstrates the feasibility of in vivo three-dimensional Compton imaging of the key ^225^Ac daughter nuclides, ^213^Bi and ^221^Fr, in tumor-bearing mice at therapeutically relevant administered activity levels (*≤* 0.5 *µ*Ci). Tumors are clearly visualized in 3D reconstructed Compton images of all three mice at tumor activity levels of approximately 30.4, 22.4, and 6.2 nCi for mouse 1, 2, and 3, respectively. To our knowledge, this is the first in vivo 3D visualization and quantification of these daughter emissions at such low injected activity. As shown in Fig. 4 and Fig. 5, both isotopes can be visualized with a single-detector scanner in a 30-min imaging session, highlighting the sensitivity advantage of our collimator-free approach. The tumor activity levels, which are on the order of tens of nCi (Table 1), indicate that the system is capable of detecting clinically relevant uptake levels for preclinical TAT studies. This capability addresses a critical limitation of conventional collimator-based SPECT systems, which typically require higher activities or extended acquisition times. An important point to notice is that our Compton imaging approach enabled tomography with a single detector view and a single bed position. This is currently not possible with standard SPECT scanners since they require several detector views and bed positions.

A key challenge in imaging ^225^Ac is the low gamma-ray yield and the complex decay scheme. Despite these constraints, the measured energy spectrum (Fig. 3(a)) and the summed-energy spectrum for *nHit* = 2 events (Fig. 3(b)) clearly show the dominant peaks at 218 keV (^221^Fr) and 440 keV (^213^Bi). These results confirm that, even at sub-*µ*Ci activity levels, sufficient event statistics can be obtained for Compton reconstruction at these two gamma energies.

An important feature of this study is the simultaneous imaging of two daughter radionuclides with gamma-ray energies of 440 and 218 keV. Notably, successful image reconstruction was achieved even at the lower energy of 218 keV through Compton imaging. A slight blurring is observed along the axial (vertical) direction as shown in the transverse and sagittal slices of Figs. 4 and 5, resulting from the single-bed-position acquisition. The ^213^Bi images exhibit superior spatial definition compared with ^221^Fr, which is attributable to the smaller angular resolution measure (ARM ~ 10^*°*^ at 440 keV versus ~ 25^*°*^ at 218 keV).

Quantitative ROI analysis supports the reliability of the imaging approach. The calibration curves (Fig. 6(a-b)) demonstrate excellent linearity between ROI counts and source activity for both energy windows (*R*^2^ *≈* 1), validating the use of linear models for activity estimation. When applied to in vivo data, the reconstructed activity values show good agreement with ex vivo biodistribution measurements (Table 1). In tumor regions, activity recovery ranges from approximately 86-98% (Fig. 7), indicating good quantitative accuracy measured by the imaging system. This level of quantitative accuracy is notable given the low counting statistics. In contrast, activity recovery values in the combined central organ region of mouse-2 exceed 100% (although within the uncertainty). This slight overestimation is likely attributable to partial-volume effects, spill-in from adjacent tissues, and the inclusion of blood-pool activity within the ROI in in-vivo imaging, whereas such contributions are excluded in ex vivo organ-specific measurements. The measured uncertainty (as summarized in Table 1), in the recovered activity for both ^213^Bi and ^221^Fr remained below 10% for all measurements except the tumor in mouse-3. The higher uncertainty of about 21% for this tumor (mouse-3) is attributable to the lower number of detected events resulting from its smaller tumor volume.

A current limitation of our method is that it is unable to spatially resolve individual organs in the central region (as seen in Figs. 4 & 5), leading to aggregation of activity in a single hot spot region. These findings suggest that quantitative Compton imaging is currently more robust for localized, high-contrast regions such as tumors than for anatomically complex regions with overlapping very small activity distributions.

Despite these limitations, the overall biodistribution in tumor and combined central organs observed in vivo are consistent with the ex vivo results (Table 1), confirming that Compton imaging can provide biologically meaningful information on radiopharmaceutical distribution. The agreement between in vivo and ex vivo measurements suggests that daughter nuclide imaging can serve as a surrogate for understanding the spatial distribution of radiation dose. Future improvements in detector design and reconstruction methodology are expected to further enhance imaging performance (*28*, *31*, *34*). Increasing detection efficiency, improving angular resolution, and incorporating attenuation and scatter corrections into the reconstruction process could enable better organ separation in the central region and reduce quantification bias. It is important to stress that this results were possible with a single detector. It is expected that a multi-detector scanner will substantially improve counting statistics and image quality, particularly for lower-energy emissions such as ^221^Fr.

From a translational perspective, the ability to perform noninvasive, longitudinal imaging of ^225^Ac daughter nuclides has significant implications. Unlike ex vivo biodistribution, which requires the sacrifice of multiple animals and provides only endpoint measurements, Compton imaging will enables repeated measurements in the same mice, allowing direct assessment of pharmacokinetics and temporal redistribution. This capability could play an important role in optimizing radiopharmaceutical design, improving dosimetry models, and ultimately enhancing the safety and efficacy of targeted alpha therapy.

## 5. Conclusion

This work establishes the capability of a collimator-free, three-dimensionally resolving CZT Compton camera to perform in vivo imaging of key ^225^Ac decay products, namely ^213^Bi and ^221^Fr, in prostate cancer mouse models under activity levels relevant to targeted alpha therapy. Using a single-detector configuration and a 30-minute acquisition, spatial distributions of these daughter radionuclides in the tumor were successfully reconstructed and coregistered with CT, enabling visualization of tumor-associated uptake as well as systemic distribution. The imaging results demonstrate that localized activity in tumors can be detected and quantified at a few tens of nCi scale, with reconstructed values showing good agreement with independent ex vivo measurements. In addition, the aggregation of signals in central body regions underscores current limitations in resolving closely spaced organs and in separating overlapping activity distributions. Despite these challenges, the overall correspondence between imaging-derived estimates and biodistribution data confirms that Compton imaging can provide meaningful and quantitative information on radiopharmaceutical behavior in vivo. With further optimization in detector configuration and reconstruction strategies, this approach has the potential to support longitudinal studies, improve dosimetric evaluation, and advance the development of targeted alpha therapeutics.

## 6. Acknowledgements

This work was supported by the NIH Grants R01EB032324 and R01CA279203. The authors thank Joshua Cates, Woon-Seng Choong, Grant Gullberg, Gyohyeok Song, Uttam Shrestha, and Robin Peter for valuable discussions. The authors also thank H3D Inc. for providing the CZT-M400 camera. The ^225^Ac was supplied by the National Isotope Development Center managed by the U.S. Department of Energy Isotope Program. The authors declare no conflicts of interest.

