## Supplemental Material for "In vivo Quantitative Tomography of ^225^Ac Daughters in a Prostate Cancer Mouse Model with a Compton Camera"

### I Approach

#### I.A Imaging system

In vivo imaging of the daughter radionuclides of $^{225}$Ac requires a highly sensitive imaging system capable of detecting extremely low activity levels. The sensitivity performance of such an imaging system was previously established and experimentally validated by our group through a dedicated detector characterization study (1). We have used a single-layered CZT Compton camera, M400, with three-dimensional interaction positioning capability (2). The detector consists of four CZT crystals, each measuring 2.2 cm×2.2 cm×1 cm ($l \times w \times t$), arranged in a planar geometry. The camera provides an energy resolution of approximately 0.67% at 440 keV and 1.23% at 218 keV (1). The interaction position resolution is approximately 0.5 mm for gamma ray energies $E_\gamma \geq 140$ keV (3). This detector enables full 3D tracking of gamma-ray events within the crystal by providing the spatial coordinates and deposited energy for each interaction. Additional technical specifications and characterization of the camera are described in Refs. (1, 2).

For the experiment, we designed a stand that holds the M400 camera in an upwards position facing the subject, which is accommodated in a cylindrical mouse bed compatible with a commercial SPECT/CT (VECTor4/CT (4)), as already illustrated in Fig. 2(a) in the main manuscript. The distance between the M400 Compton camera and the mice was approximately 5 mm. For each detected gamma-ray event, the system records the number of interactions (nHit), energy deposited ($E_i$) at each interaction site, 3D interaction coordinates ($x, y, z$), and timestamp ($t_i$). These observables are recorded in list-mode format in a binary file used for posterior analysis and image reconstruction.

#### I.B Compton imaging of $^{213}$Bi and $^{221}$Fr

Compton imaging is based on the principle that the direction of an incoming gamma ray of energy $E_0$ can be inferred if the gamma ray undergoes Compton scattering in the detector, depositing a portion of its energy $E_1$, followed by a photoelectric absorption, where the remaining energy $E_2$ is deposited. By recording the deposited energies ($E_1$ and $E_2$) and the corresponding 3D interaction positions using an imaging system, the original gamma-ray direction is constrained to a conical shell (named the

Compton cone) characterized by the Compton angle ($\theta_C$),

$$\theta_C = \cos^{-1}\left[1 - \frac{E_1 \times m_0c^2}{E_0(E_0 - E_1)}\right]; \tag{1}$$

where $m_0c^2$ is the rest mass energy of an electron (511 keV). The information contained in multiple of such conical shells is used in an image reconstruction algorithm to provide the original distribution of the gamma-ray source. A schematic representation of the Compton imaging process for in vivo mouse imaging has already been presented in Fig. 2(b) of the main manuscript.

Only events with $nHit = 2$ were selected, with a total detected energy deposition ($E_1 + E_2$) compatible with the gamma-ray energies of $^{213}$Bi ($440 \pm 3$ keV) and $^{221}$Fr ($218 \pm 3$ keV). Event selection additionally included a distance cut (dCut) of 3 mm between hits, based on our detector characterization study (1). The event selection criteria is designed to identify interactions consisting of an initial Compton scattering event followed by a subsequent photoelectric absorption, ensuring physically valid events for reconstruction and reducing contamination from incomplete energy deposition, pixel charge-sharing events, or higher-order interactions. We utilized the commonly used energy deposition comparison method to determine the sequence of two successive interactions (5–8).

##### I.C Image reconstruction

For image reconstruction, we used POSSUM (9), a Python-based Compton image reconstruction framework recently developed by our group. POSSUM incorporates a custom list-mode implementation of the Ordered Subset Expectation Maximization (OSEM, (10, 11)) algorithm tailored for Compton imaging applications. The field-of-view is discretized in a voxelized volume. The difference between the geometric angle, $\theta_{G_{ij}}$, and the Compton angle, $\theta_{C_i}$, is calculated for each event, $i$, and each voxel, $j$, as

$$\Delta\theta_{ij} = \theta_{G_{ij}} - \theta_{Ci} = \arccos\left[(\vec{S}_i - \vec{A}_i) \cdot (\vec{V}_j - \vec{S}_i)\right] - \theta_{Ci} \tag{2}$$

where $\vec{S}_i$ is the detected location of the Compton scattering for event $i$, $\vec{A}_i$ is that of the photoelectric absorption, and $\vec{V}_j$ is the location of the center of voxel $j$.

The system matrix $A$ is defined so that each element $A_{ij}$ represents the probability of an event $i$ being originated in voxel $j$. It is estimated as a Gaussian distribution for each event and voxel as

$$A_{ij} = \frac{1}{\sqrt{2\pi}\sigma} \exp -\frac{1}{2}\left(\frac{\Delta\theta_{ij}}{\sigma}\right)^2 \tag{3}$$

where $\sigma$ is a constant that represents the angular resolution. We use 10° for $^{213}$Bi and 25° for $^{221}$Fr, as calculated in Ref.(1).

The system matrix is stored in memory as a sparse matrix by only recording those values above 10% of the maximum probability for each event. The system matrix is generated using GPU computation and stored in CPU memory for subsequent use.

The full dataset of Compton events, $N$, is divided in subsets, each containing $N_s$ events with a corresponding subset of the system matrix $A^s$. Then, the OSEM iterative formula is applied

$$\lambda_j^{k+1} = \frac{\lambda_j^k}{\eta_j} \sum_i^{N_s} \frac{A_{ij}^s}{\sum_l A_{lj}^s \lambda_j^k} \tag{4}$$

where, $\lambda_j^k$ and $\lambda_k^{k+1}$ are the activity in the $j^{th}$ voxel at iteration steps $k$ and $k+1$, respectively, and $\eta_i$ denotes the sensitivity associated to the $j^{th}$ voxel. This formula is applied to each subset $A_s$ in succession using GPU, which completes a full iteration. In this study, we use 50 iterations and one subset as optimized.

#### I.D Region-of-interest (ROI) calibration

For quantitative analysis of the in vivo Compton images, calibration measurements were performed using $^{225}$Ac point sources with low-activities of 7.8, 11.4, 17.8, 33.4, and 57.3 nCi. Each dataset was acquired for 30 minutes under identical experimental conditions. Three-dimensional Compton images of the daughter nuclides $^{213}$Bi and $^{221}$Fr were reconstructed for each source using the same event selection criteria and reconstruction parameters as applied to the in vivo imaging data, ensuring consistency between calibration and animal studies. Quantitative regions of interest (ROIs) calibration of the Compton images was performed using the Medical Imaging Data Examiner (AMIDE) software. The ROIs were defined around the reconstructed point sources, and the total counts within each ROI were extracted. The ROI counts corresponding to the 440 keV ($^{213}$Bi) and 218 keV ($^{221}$Fr) gamma energies were plotted as a function of source activity and subsequently fitted using a linear model,

$$y(x) = ax + b, \tag{5}$$

where $y(x)$ represents the measured ROI counts at activity $x$, $a$ is the sensitivity factor, and $b$ accounts for background and noise contributions. The resulting calibration parameters ($a$ and $b$) were obtained from these fits.

For in vivo image quantification, three-dimensional ROIs were delineated around the tumor and the central activity regions in both the reconstructed $^{213}$Bi and $^{221}$Fr images to obtain the corresponding ROI counts for all mice. These ROI counts from the in vivo mouse images were subsequently converted into activity uptake for $^{213}$Bi and $^{221}$Fr using the calibration parameters ($a$ and $b$) derived from the ROI calibration analysis.

#### I.E Ex vivo bio-distribution measurement

Immediately after completion of the in vivo imaging session on day 7 p.i., all animals were euthanized to perform ex vivo biodistribution measurements for direct comparison and validation of the quantitative in vivo imaging results. Tumors and major organs, including the kidneys, liver, heart, lungs, brain, spleen, pancreas, large intestine, small intestine, and stomach, were excised and collected. In addition, samples of muscle, bone, and blood were obtained. The radioactivity in each specimen was measured using a HIDEX gamma counter to quantify organ-specific activity distribution. The gamma-ray energy spectrum and mass of each specimen were measured. The activity concentrations and the activity uptake were determined for all samples across the three mice based on selected gamma-ray peaks, 440 keV for $^{213}$Bi and 218 keV for $^{221}$Fr.

##### I.F Activity recovery

We quantitatively evaluate the accuracy of the in vivo images by comparing the activity estimates derived through the ROI imaging analysis against the current gold standard quantification via ex vivo biodistribution. The activity recovery ($R$) was calculated as

$$R = \left[1 - \frac{A_{\text{ex vivo}} - A_{\text{in vivo}}}{A_{\text{ex vivo}}}\right] \times 100\%, \tag{6}$$

where $A_{\text{ex vivo}}$ and $A_{\text{in vivo}}$ represent the activities obtained from ex vivo measurements and in vivo ROI-based quantification, respectively.

#### II Results

##### II.A Activity concentration of tumor and the central organs

Table 1: Activity concentrations in the tumor and combined central organ region were obtained from the ex vivo measurement using HIDEX.

| | Organ name | Weight (gm) | [a]Activity Concentration (%/gm) | |
|---|---|---|---|---|
| | | | **$^{213}$Bi** | **$^{221}$Fr** |
| **Mouse-1** | Tumor | 0.41 | 15.4±0.3 | 16.2±0.4 |
| | Central organs | 1.27 | 2.8±0.1 | 2.4±0.1 |
| **Mouse-2** | Tumor | 0.23 | 17.3±0.4 | 18.4±0.4 |
| | Central organs | 1.90 | 2.0±0.1 | 1.9±0.1 |
| **Mouse-3** | Tumor | 0.11 | 13.3±0.5 | 16.4±0.6 |
| | Central organs | 2.08 | 8.7±0.2 | 8.4±0.2 |

[a]The organ's uptake, activity per gram (activity concentration in %/gm) w.r.t. total activity obtained from the ex vivo biodistribution measurement.
Central organs include the heart, liver, spleen, kidneys, and lungs.
